# Molecular Insights into Single Chain Lipid Modulation of Acid-Sensing Ion Channel 3

**DOI:** 10.1101/2024.08.29.610156

**Authors:** Ramya Bandarupalli, Rebecca Roth, Robert C Klipp, John R Bankston, Jing Li

**Affiliations:** Department of Biomolecular Sciences, School of Pharmacy, University of Mississippi, Oxford, MS; Department of Physiology and Biophysics, University of Colorado Anschutz Medical Campus, Aurora, CO

## Abstract

Polyunsaturated fatty acids (PUFAs) and their analogs play a significant role in modulating the activity of diverse ion channels, and recent studies show that these lipids potentiate acid-sensing ion channels (ASICs), leading to increased activity. The potentiation of the channel stems from multiple gating changes, but the exact mechanism of these effects remains uncertain. We posit a mechanistic explanation for one of these changes in channel function, the increase in the maximal current, by applying a combination of electrophysiology and all-atom molecular dynamics simulations on the open-state hASIC3. Microsecond-scale simulations were performed on open-state hASIC3 in the absence and presence of a PUFA, docosahexaenoic acid (DHA), and a PUFA analog, N-arachidonyl glycine (AG). Intriguingly, our simulations in the absence of PUFA or PUFA analogs reveal that a tail from the membrane phospholipid POPC inserts itself into the pore of the channel through lateral fenestrations on the sides of the transmembrane segments, obstructing ion permeation through the channel. The binding of either DHA or AG prevented POPC from accessing the pore in our simulations, relieving the block of ionic conduction by phospholipids. Finally, we use the single-channel recording to show that DHA increases the amplitude of the single-channel currents in ASIC3, which is consistent with our hypothesis that PUFAs relieve the pore block of the channel induced by POPCs. Together, these findings offer a potential mechanistic explanation of how PUFAs modulate ASIC maximal current, revealing a novel mechanism of action for PUFA-induced modulation of ion channels.

## Introduction

Acid-sensing ion channels (ASICs) are ligand-gated sodium channels found in the nervous system that respond to alterations in extracellular pH, playing a pivotal role in signal transduction associated with pH changes (1). While primarily implicated in pain perception and inflammation in the peripheral nervous system (PNS), ASICs also have important implications in the central nervous system (CNS), contributing to fear conditioning, ischemic cell death, and synaptic plasticity(1–5).

ASICs can exist as both homotrimers and heterotrimers, wherein each subunit consists of two transmembrane helices, intracellular N- and C-termini, and a bulky extracellular domain. ASICs are activated by rapid acidification, which is thought to protonate critical residues in the extracellular domain leading to a conformational change that allows the channel to transition from a resting state to an open state and then to a desensitized state (5,6). During acidification, the transmembrane helices that comprise the pore region exhibit significant rearrangement between the conducting (open) state and the non-conducting (desensitized and resting) states, which is consistent with the opening/closure of the pore (2,3,7–10).

Structures of ASIC1 from chicken also revealed lateral fenestrations near the extracellular vestibule, which are thought to act as a secondary pathway for ion permeation (6,11,12). These fenestrations start in the extracellular space but extend significantly into the lipid bilayer. Upon transitioning between conducting and non-conducting states, the rearrangement in the extracellular and transmembrane domains allows for these lateral fenestrations to expand, and the openings on the extracellular side have been suggested to provide access to the pore for both ions and pore blockers such as amiloride (11). However, the role of the fenestrations within the bilayer remains unknown.

ASIC3 is one of 7 mammalian subunits encoded by 5 ASIC genes (ACCN 1-5) and the most sensitive isoform to changes in extracellular pH (1,3,5). ASIC3 is expressed in many tissues but has been primarily studied in the PNS where they have been shown to play a role in acid-induced cutaneous pain, itch, hyperalgesia, and inflammatory pain (13–17). Additionally, ASIC3 activity can also be modulated by proalgesic single-chain acyl lipids that are readily released from the membrane following inflammation (14). The concentration of proalgesic lipids such as arachidonic acid (AA) and lysophosphatidylcholine (LPC) was shown to be increased in exudates from patients with inflamed joints and could even induce ASIC3 activation in the absence of changes in pH(14). Moreover, LPC and AA alone were enough to increase pain sensitivity in an ASIC3-dependent manner when injected into the hindpaws of mice (14).

Polyunsaturated fatty acids (PUFAs) such as AA can alter the function of a number of ion channels including Na^+^, Ca^2+^, and K^+^ ion channels (18–21). Recently, we have shown that many PUFAs and PUFA analogs, such as N-acyl amino acids (NAAAs) like N-arachidonyl glycine (AG), potentiate ASIC3 and cause multiple functional changes in the channel that lead to an increase in the overall current (18). First, lipid binding to the channel causes a shift in the pH dependence of activation in the basic direction, allowing for the channels to activate under more physiological pH conditions. In some cases, the magnitude of this shift is sufficient to cause channel opening even at neutral pH. Second, lipid interaction leads to a slowing of channel desensitization. Finally, these lipids increase the magnitude of the maximal ASIC3 current (I^Max^) at saturating concentrations of protons. Moreover, an arginine, (R63 in human ASIC3 (hASIC3), R64 in rat ASIC3), located in the outer leaflet of TM1 was identified to be a critical residue for PUFA potentiation in ASIC3 (18). A recent study has also suggested that AA makes several coordinating interactions between this arginine along other ASIC3 TM residues within the lower region of the fenestrations (22). Therefore, it is possible that these fenestrations could potentially serve as a universal binding region for various classes of lipids including PUFAs, PUFA analogs, and native phospholipids within the membrane.

Despite the potential importance of these lipids in influencing inflammatory pain signaling via ASIC3, we do not yet have a clear atomistic or mechanistic framework for understanding their effects. Here we have used microsecond-scale unbiased molecular dynamics (MD) simulations and patch-clamp electrophysiology to reveal a potential mechanism for the change in I_Max_ that occurs upon lipid binding. We compared simulations in the absence and presence of AG or DHA and found that both potentiating lipids alter the interactions between the channel and the native phospholipids in a manner that may change the gating of the channel.

## Materials and methods

### Homology Modelling & Molecular Docking

In the absence of experimentally resolved structures for human ASIC3 on the Protein Data Bank (PDB), homology modeling was utilized to model its structure in the open state. This study employed the existing chicken ASIC1 (PDB ID: 4NTY) structure in the open state as templates for homology modeling (23). The protein sequence of hASIC3 was obtained from the UniProt database (UniProt ID: Q9UHC3 sharing 49.71% sequence identity with chicken ASIC1) and subsequently submitted to the Swiss-Model web server (version 1.2) (24,25). SwissModel generated 12 structure models for the open state of hASIC3. The selection of the most suitable model was based on the Global Model Quality Estimate (GMQE) scores (26), which evaluate the overall quality of the model. The structure with the highest GMQE score (0.68) was chosen for the open state of hASIC3. This selected model was then utilized to perform molecular docking and build simulation systems for molecular dynamics (MD) runs.

Molecular docking was performed in Schrodinger’s suite (27). From the protein preparation wizard, the protein was pre-processed by assigning bond orders, adding hydrogens, ensuring proper ionization at pH 7.4, creating disulfide bonds, and deleting waters. The protein structure is optimized for the H-bond assignment and minimized. The 3D structure of DHA was obtained from PubChem, whereas the 3D structure of AG was built in Maestro and minimized. 3D structures of AG & DHA are subject to Ligprep (28), in which ligand possible states between target pH 8.0 to 6.0 were generated and the chirality from the 3D structure was determined. An arginine, R63, located in the outer leaflet of transmembrane domain 1 (TM1) was identified to be a critical residue for PUFA potentiation in rat ASIC3 (18). Hence, the receptor grid is generated with R63 as the centroid. For the ligand midpoint box, the box length in the X, Y, and Z axes was set at 30, 30, and 30 Å respectively. Following receptor grid generation, molecular docking was performed using the Glide function of Maestro (29,30). For the docking, Extra precision docking (XP) mode was used (31). Finally, the interactions of selected ligand and protein-docked complexes were analyzed by the pose viewer. The structures of the best pose with the highest G-scores were used for MD simulations. Since ASIC3 is a homotrimer, it has 3 similar lipid binding sites, hence, 3 PUFA (3-AG & 3-DHA) molecules were docked to R63 of open-state ASIC3 and simulated to uncover the modulation mechanisms.

### Molecular Dynamics Simulation system settings

#### System setup

All the simulation systems were built in CHARMM-GUI Membrane Builder (32). Disulfide bonds were added between the cysteine residue pairs (92-186, 164-171, 282-370, 315-366, 319-364, 328-350, 330-342) to ensure the structural integrity and stability of the protein. To determine the appropriate protonation states of ionizable residues, pKa calculations were performed using PROPKA3 (33). Based on the obtained pKa values all ionizable residues were assigned the default protonation states in the simulation system. The protein was then embedded into the lipid layer (Table 1) and the systems were solvated in a 0.15M NaCl aqueous solution using the web service CHARMM-GUI (34).

**Table 1:**
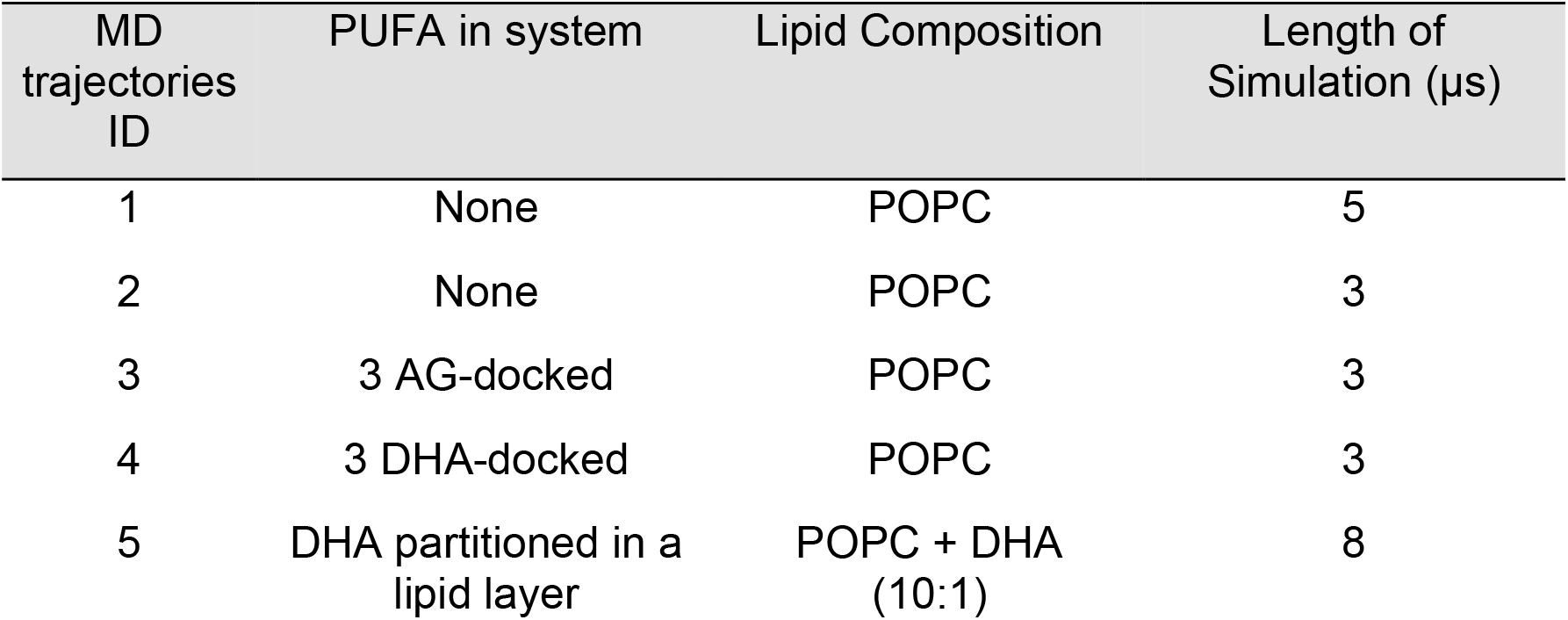
Summary of simulation systems.

| MD trajectories ID | PUFA in system | Lipid Composition | Length of Simulation (μs) |
| --- | --- | --- | --- |
| 1 | None | POPC | 5 |
| 2 | None | POPC | 3 |
| 3 | 3 AG-docked | POPC | 3 |
| 4 | 3 DHA-docked | POPC | 3 |
| 5 | DHA partitioned in a lipid layer | POPC + DHA (10:1) | 8 |

For trajectory (Traj.) 1&2, the open-state hASIC3 protein was embedded within the phosphatidylcholine (POPC) bilayer in the absence of PUFAs, which hereafter are referred to as the “apo” form. PUFA-bound structures obtained from molecular docking of 3-AG and 3-DHA were simulated in Traj. 3 & 4 respectively. For Traj.5, the lipid composition of POPC:DHA in a 10:1 ratio with DHA molecules randomly partitioned into the lipid layer. Notably, the concentration of DHA in this configuration exceeded physiological membrane compositions. However, this intentional increase facilitates a comprehensive and efficient exploration of contacts between DHA and hASIC3, enabling reproducible measurements of lipid occupancy and interaction duration without compromising membrane integrity.

#### All-atom MD Simulation protocol

All-atom MD simulations were performed using either NAMD 2.14 (35) or the specialized computational platform Anton2 (36). The CHARMM36m force field was used for proteins, lipids, and ions within the study (37–39). The explicit representation of water molecules was achieved through the implementation of the TIP3P model (40). The simulations were executed under NPT (constant number of particle N, pressure P, and temperature T) conditions at a temperature of 310 K and a pressure of 1 atm, utilizing periodic boundary conditions. Electrostatic interactions were handled using the particle mesh Ewald method with a 12 Å cutoff for real-space interactions (41). Hydrogen atom bond distances were constrained through the application of the SHAKE algorithm (42). Hydrogen mass repartitioning (HMR) allows us to re-distribute the mass of heavy atoms to the bonded hydrogens (43). Hence a time step of 4 fs was selected for the simulations ran on NAMD. In MD simulations conducted with Desmond on Anton2, a Berendsen coupling scheme was implemented to sustain a consistent pressure of 1.0 atm. The calculation of long-range electrostatic interactions was facilitated by the k-space Gaussian split Ewald method (44), utilizing a 64 × 64 × 64 grid. Time-step for simulations run on Anton2 is 2.5 fs. Prior to the production runs, minimization and equilibrations were conducted for 2ns with harmonic positional restraints. The equilibrated systems were simulated at microseconds timescale. MD trajectories were then visualized and analyzed using VMD (45), in-house Tcl, and Python scripts.

#### Water occupancy analysis

For the assessment of pore blocking, the quantification of water molecules within the channel pore was performed as a metric. In the analysis of water permeation, the entirety of the channel pore was discretized into 15 cylindrical sections within the X & Y area of 120 Å^2^ and a 1Å segment along the Z-axis. The calculation involved determining the number of water molecules present within each of these defined sections with the function of time. This spatial partitioning strategy facilitated a detailed examination of water permeability through the channel pore.

#### Occupancy volume map of lipids

To understand the occupancy of POPC/PUFA lipids in the lipid layer, especially in the vicinity of the channel, we used the VolMap plug-in from VMD 1.9.4 for the analysis (45). Grid resolution was set at 1Å and the 2-D occupancy maps were generated for the entire POPC molecules. A scale value of 0.3 signifies that the PUFA was engaged with the respective residue for more than 30% of the simulation duration.

#### Ion permeation analysis

To assess the ion permeation events through the pore along the trajectories, we tracked the z-position of all the ions that enter the transmembrane domain pore during MD simulations. Ion permeation events would be indicated by a complete trace across the transmembrane pore. In addition to water occupancy, the lack of ion permeation events during a certain period of simulation time further demonstrates the pore’s impermeability.

#### Materials and Plasmids

Docosahexaenoic acid (DHA) and N-arachidonyl glycine (AG) used in electrophysiology experiments were purchased through Cayman Chemical. ASIC3 plasmid from rats was gifted by David Julius (University of California, San Francisco, San Francisco, CA) and subcloned into a pcDNA3.1 vector. To visualize cell expression, the fluorescent tag mCerulean3 was attached to the C-terminus of the channel using a short proline-rich linker, which has been previously reported to have minimal effects on channel gating (18).

#### Cell culture and transfection

CHO-K1 cells (ATCC) were cultured using Ham’s F12 medium with 10% FBS at 37°C in 5% CO_2_. Cells were grown to ∼70-80% confluency and were transfected with rat ASIC3 plasmid DNA via electroporation with a Lonza 4D Nucleofector unit following the manufacturer’s protocols. Following transfection, cells were plated on 12-mm glass coverslips coated in poly-L-lysine and incubated at 37°C in 5% CO_2_.

#### Preparation and application of lipids

Electrophysiological experiments in the presence of DHA and AG were performed under identical conditions as control experiments except solutions contained the indicated concentration of lipids. Lipid stock solutions were made up of ethanol added to aqueous solution following the manufacturer’s recommendations. Ethanol solvent in the final solution was typically 0.01% and never exceeded 0.1%. Solution pH was measured prior to and after the addition of PUFAs to the solution to ensure no pH change occurred.

#### Electrophysiological recordings

As previously described (18), whole-cell recordings were performed 16–30 h after transfection. To assess peak current magnitudes, whole-cell patch-clamp configuration was used. Borosilicate glass pipettes (Harvard Apparatus) were pulled to a resistance of 2–6MΩ (P-1000; Sutter Instrument) for whole-cell experiments. Glass pipettes were filled with an internal solution containing (in mM) 20 EGTA, 10 HEPES, 50 CsCl, 10 NaCl, and 60 CsF, pH 7.2. The extracellular solution contained (in mM) 110 NaCl, 5 KCl, 40 NMDG, 10 MES, 10 HEPES, 5 glucose, 10 Trizma base, 2 CaCl2, and 1 MgCl2, and pH was adjusted as desired with HCl or NaOH. An Axopatch 200B amplifier and pCLAMP 10.6 (Molecular Devices) were used to record whole-cell and single-channel currents. Recordings were performed at a holding potential of −80 mV with a 5-kHz low-pass filter and sampling at 10 kHz. Solution changes were performed through rapid perfusion using an SF-77B Fast-Step perfusion system (Warner Instruments). Fluorescence was visualized on an Olympus IX73 microscope with a CoolLED pE-4000 illumination system.

To measure the increase in I_Max_, cells in the absence of PUFAs were exposed to a holding pH of 8 for 6s followed by a 1.5-s application of pH 5.5. The activation protocol was then repeated with 20μM DHA or AG in both pH 8 and pH 5.5 solutions. The pH 5.5 current peaks in the presence of lipids were divided by the pH 5.5 currents in the absence of lipids to determine the percentage increase of Imax. Each control experiment was measured for five sweeps and then averaged.

For single-channel experiments, recordings were made from excised patches in the outside-out patch clamp configuration with pipettes pulled to a resistance of 6-10MΩ. Solutions were the same as in the whole cell experiments except that the extracellular NaCl concentration was increased from 110 to 150mM to increase current magnitudes. Currents were filtered using the four-pole Bessel filter of the Axopatch 200B at 2kHz and filtered digitally in the Clampex software (Molecular Devices) down to 1kHz. Patches with a steady baseline, less than 1pA leak, and fewer than 5 simultaneous openings were used for analysis. An all-points current amplitude histogram was generated using Clampex. We then fit all single opening events to a Gaussian in MATLAB (Mathworks).

#### Statistics

All functional data are plotted as the mean +/-the standard error of the mean.

Appropriate statistical tests are used and described in the figure legends.

## Result

### AG and DHA potentiate ASIC3 current in pH-saturating whole-cell recordings

We previously showed that both NAAAs and PUFAs increase ASIC3 current even at saturating proton concentrations. To show how lipids alter I_Max_ in ASIC3 we tested two lipids for their effect on ASIC3: N-arachidonyl glycine (AG) and docosahexaenoic acid (DHA). While both are single acyl chain lipids, they have different head groups, tail lengths, and double bond positions (Fig. 1A). To measure the change in I_Max_ for ASIC3, we recorded currents from Chinese Hamster Ovary (CHO) cells expressing rat ASIC3 (rASIC3) with a C-terminal Cerulean tag in the whole cell patch clamp configuration. The half-maximal activating pH (pH_0.5_) of rASIC3 varies slightly from study to study, but in all cases, the curve is fully saturated at pH 5.5 (9,18,46). To elicit rASIC3 currents, we rapidly switched solutions from pH 8 to pH 5.5 for 1.5 s and then returned to pH 8 for 6 s. We repeated this five times to ensure our currents were stable and then switched to a pH 8 solution that contains 20μM DHA and repeated this measurement for up to 90 s. Figure 1B shows representative currents using this protocol, and it is apparent that the whole cell current amplitude increases as DHA is washed on. We previously found that this wash on was mostly saturated after one minute (18). Examining this effect across multiple recordings, we found that there was an initial increase in I_Max_ of about 16% +/-4% after 3 s of DHA exposure, and after prolonged exposure (85.5 s) the increase in I_Max_ stabilized to 34% +/-5%. We then repeated this same experiment to assess the effect of 20 μM AG on I_Max_. The AG effect saturates much more rapidly (18) and here showed a larger increase in I_Max_ of about 67% +/-10% (Fig. 1C). Despite the large change in total current, there have been no mechanistic examinations of how these lipids alter ASIC3. Therefore, we utilized MD simulations to investigate a potential mechanism that aligns with the observed increase in I_Max_.

**Figure 1.**
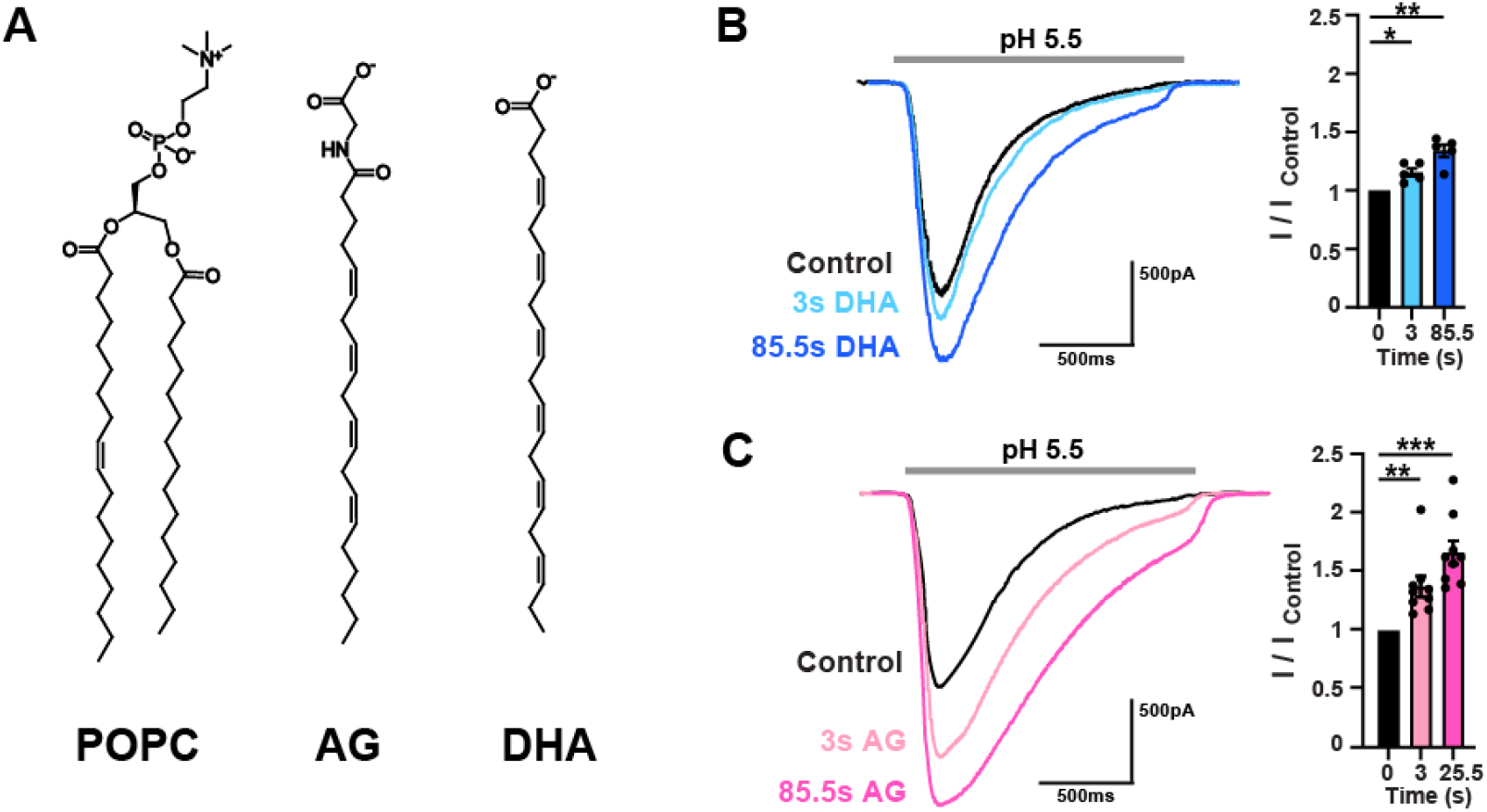
DHA and AG increase the maximal current of ASIC3 in CHO-K1 cells. **(A)** Structures of POPC (left), AG (center), and DHA (right). **(B)** Left: Representative whole-cell recordings showing pH 5.5-evoked ASIC3 currents from a single cell at different exposure times to 20µM DHA. Right: Bar plot showing the fractional change in the maximal current measured at pH 5.5 (I_max_) in the absence of DHA (black), and in response to DHA at 3 seconds (light blue; fold-increase = 1.16 ± 0.04, *n* = 5 cells) and 86 seconds (dark blue; fold-increase = 1.34 ± 0.05, *n* = 5 cells) exposure times. **(C)** Left: Representative whole-cell recordings showing 5.5-evoked ASIC3 currents from a single cell at different exposure times to 20µM AG. Right: Bar plot showing the fractional change in the maximal current measured at pH 5.5 (I_max_) in the absence of AG (black), and in response to AG at 3 seconds (light pink; fold-increase = 1.38 ± 0.09, *n* = 9 cells) and 26 seconds (dark pink; fold-increase = 1.67 ± 0.1, *n* = 9 cells) exposure times. All data given as mean ± SEM, between-group comparisons as determined by a one-way ANOVA with a post hoc Dunnett’s tests. *, P < 0.05; **, P < 0.01; ***, P < 0.005 (see Materials and methods for details).

### Tails of POPC lipids enter and block the central pore of hASIC3

Our long-term goal is to understand how lipids interact with and regulate ASIC3 function. To begin to uncover mechanisms of lipid regulation of ASIC3, we performed multiple µs-scale all-atom MD simulations (Table 1) using a homology model of an open human ASIC3 (hASIC3) structure embedded in POPC lipid bilayer in the presence and absence of a single acyl chain lipids AG and DHA (Fig. 2A, see methods). In our apo simulations, which only contain POPCs in the lipid bilayer, we consistently observed that the tails of the POPC from the upper leaflet of the membrane penetrated the pore of the channel through its lateral fenestrations (Fig. 2A). This phenomenon can be seen in two independent simulations, which we call trajectory 1 (Traj.1) and trajectory 2 (Traj. 2).

**Figure 2.**
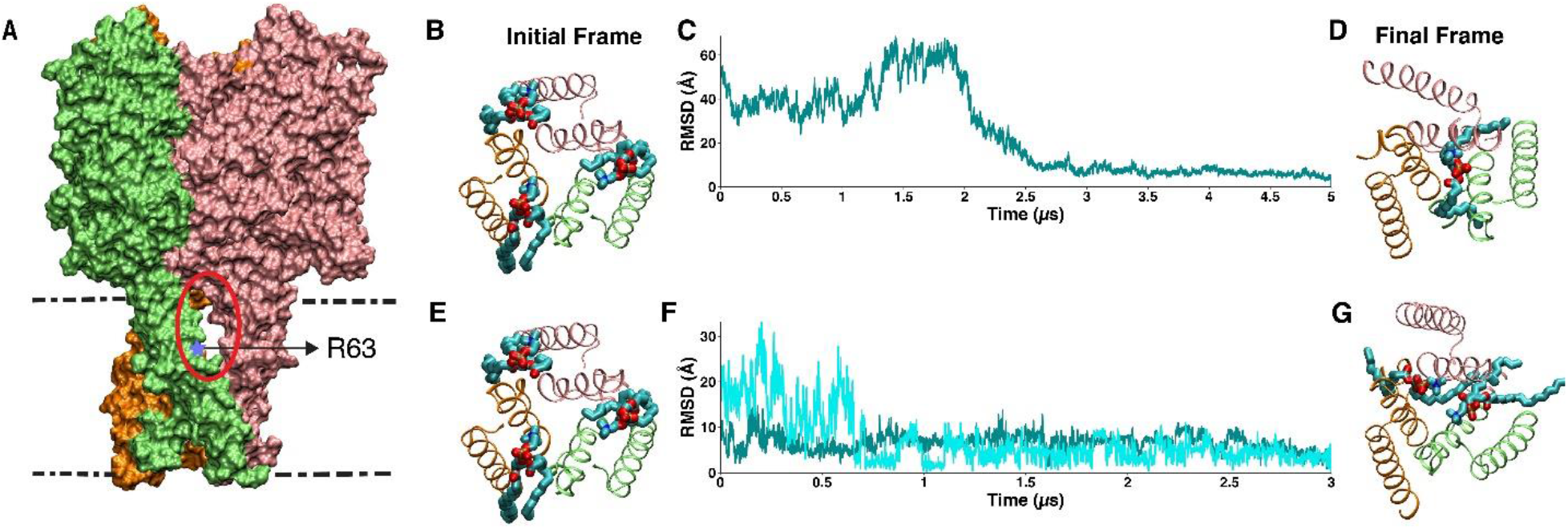
POPC tails enter the pore of open-state hASIC3. **A)** Homology model of the open state structure of hASIC3 used throughout simulations. The lateral fenestration site is circled in red and the arginine residue found to be critical for DHA potentiation is indicated with a blue star. **B)** Top view of the first snapshot of Traj.1. POPC entry into the pore did not occur initially at the beginning of the simulation. **C)** RMSD plot of POPC in Traj. 1: POPC entered the pore at around 2.5 µs and stayed in the pore for the remainder of the simulation. **D)** Top view of the last snapshot of Traj. 1, which was used as a reference point to measure the RMSD of POPC throughout the simulation. **E)** Top molecular view of the first snapshot of Traj.2. POPC entry into the pore did not occur initially in the simulation. **F)** RMSD plots of two POPCs in Traj. 2: the 16:0 tail from a POPC lipid entered the pore within the first 500ns (teal), and an additional 18:1 tail from another POPC lipid penetrated the pore around 600ns (cyan). **G)** Top molecular view of the last snapshot of Traj. 2, which was used as a reference point to trace the movement of POPCs throughout the simulation.

To examine the dynamics of the lipid penetration into the pore, we monitored the root mean squared displacement (RMSD) of the POPC that entered the pore across the whole trajectory (Fig. 2C, F) using its position in the last frame as the reference (Fig. 2D, G). In the initial conformation of Traj. 1, the pore appeared open without any block from the membrane lipids (Fig. 2B). However, between 2 and 2.5 µs the 16:0 lipid tail from a POPC entered the pore through the fenestrations of the channel where it resided for the duration of the simulation. The whole process can be observed from the drop in the RMSD value from 50 Å at the beginning of the simulation to about 10 Å at 2.5 µs where it remained until the final frame (Fig. 2C, 2D). In Traj. 2, we observed two different lipid tails entering the pore through the fenestration. Again, the pore was open at the start of our simulation (Fig. 2E). Then, at ∼200ns a POPC lipid-tail (16:0) penetrated the pore, as shown by the RMSD value dropping 10 Å relative to the final frame (Fig. 2F). Additionally, we also observed the 18:1 tail from a second POPC accessing the central pore at around 600 ns, resulting in two lipid tails simultaneously occupying the pore (Fig. 2F&G). Although hASIC3 interacted with many POPC lipids in the simulated bilayer, not all POPC molecules in the vicinity of the fenestration site access the central pore. The POPC lipid initially closest to the fenestration site, along with many other POPC lipids bound to the fenestration site during the simulations, did not enter the pore (Fig. S1). In contrast, the POPC lipids that penetrated the central pore diffused towards R63 (Fig. 2A) from more distant locations, as shown by their initial RMSD values (30 to 50 Å) (Video S4). This suggests that the POPC lipids bound to the fenestration site do not always block the central pore, indicating a stochastic process with a certain probability.

Next, we visualized the blockade of the pore by plotting occupancy volume maps of POPC. These maps provide a visual for where in space POPC resides throughout the entire simulation. Figure 3A shows a top-down view of hASIC3 with a heatmap that color codes the fractional occupancy of POPC in the XY panel. The blue regions, where there is no POPC, is the volume that hASIC3 occupies. The molecular view of these heatmaps can be seen in Figure 3B. In both trajectories, POPC surrounded the channel and, interestingly, showed high POPC occupancy in the pore of the channel (Fig. 3A). By entering the pore through the lateral fenestration, the lipid tails interrupted the water pathway of the pore through which ions translocate. Therefore, pore blockade was quantitatively assessed through water occupancy analysis. In short, this approach allows us to determine if there is an aqueous pathway through the channel pore by determining where water molecules are present in each snapshot of our simulations. If POPC lipid tails occlude the pore, the aqueous path would be interrupted and permeation reduced or even eliminated. For both trajectories, the number of water molecules within the pore along the Z-axis was monitored throughout the entire simulation (Fig. 3C). We converted this data into heat maps that show the amount of water along the length of the pore. In Traj. 1, there were consistently between 1-4 or more water molecules all through the pore for the first 2.5 µs. This aqueous path was then interrupted once the POPC tail entered the pore. In Traj. 2, we saw a similar phenomenon where the aqueous pathway through the channel was disrupted throughout much of the simulation, although the dehydration region was not as wide as that in Traj.1 (Fig. 3C). This occlusion of the water pathway can be visualized in a structural image, demonstrating where water was absent from the central portion of the permeation path (Fig. 3D).

**Figure 3:**
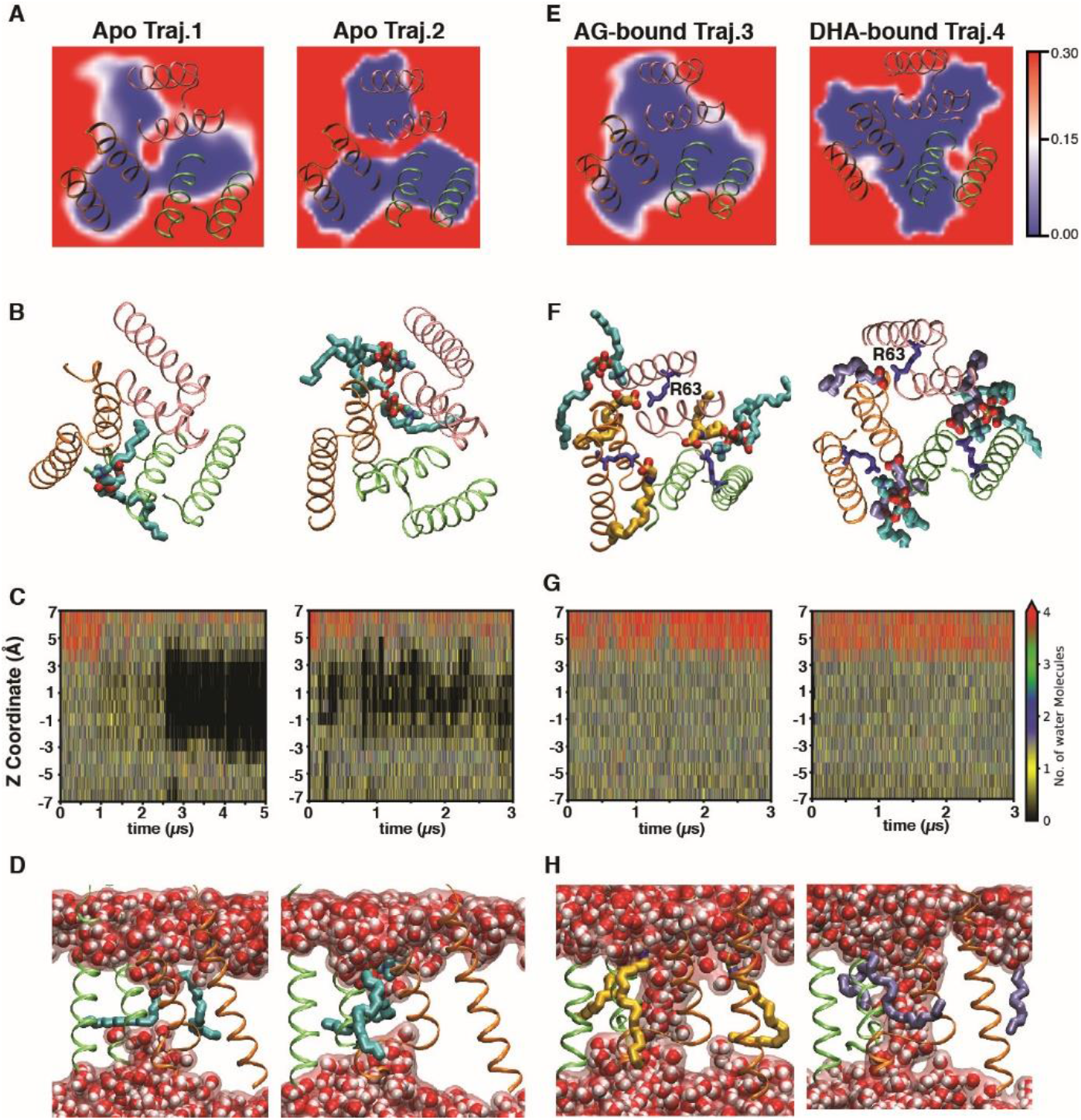
AG and DHA prevent POPC from blocking the pore and occluding water permeation through hASIC3 in the open state. **A)** Occupancy volume maps of POPC. The color bar represents the fractional occupancy of POPC, where 0.3 corresponds to at least 30% occupancy during the simulation. High occupancy can be seen in the central pore of hASIC3 in both Traj. 1 and Traj. 2. **B)** Molecular images of the top view of POPC (represented in cyan licorice) in the pore in Traj.1 and Traj.2. Snapshots are obtained from the last frame of trajectories. **C)** Water permeation heatmaps, which show the number of water molecules along the Z-coordinate (Y-axis) over time (X-axis). The scale bar shows that anywhere from 0-4 or more water molecules can be present along the Z-coordinate of the pore region. Black regions in this map indicate dehydrated areas within the pore at specific time points. **D)** Molecular images of the last frame of Traj. 1&2 to show POPC interrupting water wire. **E)** Occupancy volume maps of POPC in simulations with AG-bound (Traj. 3) and DHA-bound (Traj. 4) ASIC3, presented in a similar manner to Fig. 3A. **F)** Molecular images of the top view of POPC and AG (yellow licorice) or DHA (ice blue licorice), which both prevent POPC entry to the pore in the representative snapshots. **G)** Water permeation heatmaps for AG-bound (Traj. 3) and DHA-bound (Traj. 4) ASIC3, presented in a similar manner to Fig.3C. **H)** Molecular images of continuous water flowing through the AG-bound (Traj. 3) and DHA-bound (Traj. 4) ASIC3 pore.

### AG and DHA binding prevents phospholipid block of the hASIC3 central pore

Based on these *in silico* observations from the apo simulations, we hypothesized that the increase in I_Max_ in the presence of potentiating lipids might reflect a relief of this block resulting from the displacement of POPC lipids from the binding region on the channel and thereby preventing tail occlusion of the pore. Our previous work showed that PUFAs and PUFA analogs act via the outer leaflet of the membrane (18). As noted above, we found that an arginine located within the fenestration site, R63, was necessary for DHA to alter ASIC3 function. Mutation of this residue to glutamine eliminated the shift in the pH dependence of activation, the slowing of the desensitization rate, and the change in I_Max_ (18). This suggests the possibility that this arginine is a critical interacting residue for PUFAs.

Based on this experimental data, we designed our PUFA-bound simulations where we docked either AG (AG-bound Traj. 3) or DHA (DHA-bound Traj. 4) to all three equivalent sites (the R63 site) in the open state of hASIC3. We then performed 3 µs simulations of both the 3-AG and 3-DHA bound form of hASIC3. In both simulations, AG and DHA remained bound to the R63 site for the entire simulation (Fig. S2). Throughout both the AG- and DHA-bound trajectories, no penetration of POPC lipid tails into the pore was observed (Video S5). This can be seen by the absence of POPC in the central pore in occupancy volume maps (Traj. 3 & 4, Fig. 3E) as well as in the structural images (Fig. 3F). This relief of block also predictably resulted in the continuous presence of water in the pore (Fig. 3G, H), suggesting the aqueous path was open in the presence of DHA or AG.

### AG and DHA restoration of the water pathway allows for Na^+^ conduction

Alterations to the central pore’s aqueous pathway would likely disrupt the ability of ASIC3 to permeate Na^+^ ions through the pore. To assess how disrupting the water path would impact ion flux, we performed an ion permeation analysis where we tracked the z-position of all ions that entered the transmembrane region of the channel during our simulations. Figures 4A & B show the position of ions throughout the simulation for both apo trajectories. In Traj.1, several permeation events can be seen where ions cross from one side of the channel to the other. However, after ∼2 µs, when the POPC tail entered the pore there were no ions observed translocating through the pore for the rest of the simulation (Fig. 4A). Similarly, for Traj. 2, a few ions passed through the channel at points where the tail block is incomplete and the aqueous path is open, but throughout the rest of the simulation, no ions traversed the pore (Fig. 4B). This confirms that the central pore cannot permeate Na^+^ after POPC lipids enter the central pore in Traj. 2, even though the dehydration of its pore is less than that of Traj. 1.

**Figure 4:**
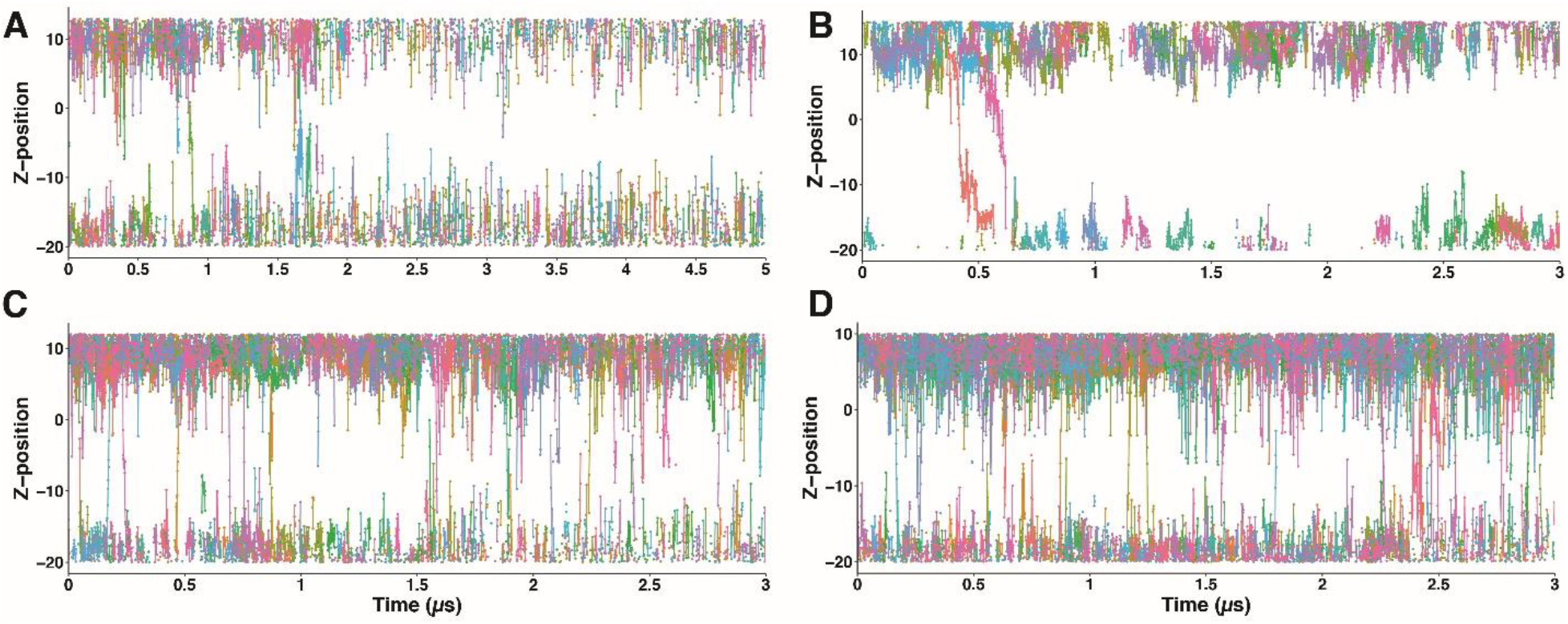
Na^+^ ion permeation through the hASIC3 pore in apo, AG-bound, and DHA-bound states. (A & B) Apo state (Trajs. 1 & 2). (C) AG-bound state (Traj. 3). (D) DHA-bound state (Traj. 4). Each plot shows the Z-position of Na^+^ ions (Y-axis) over time (X-axis), with individual ions represented by different colors. In the apo form, ion permeations occur primarily before POPC entry. In contrast, AG-bound and DHA-bound states exhibit ion permeation events throughout the entire simulation duration.

AG and DHA both precluded the blockage of the pore by POPC tails through binding to the critical arginine R63 preventing POPC access to the fenestration region of the channel. We found that this open pathway in AG- and DHA-bound trajectories also allowed for the restoration of ion permeation through the pore (Fig.4 C & D). Upon AG or DHA binding, ion permeation events are observed through the whole trajectories. This observation suggests that the binding of PUFA or PUFA analog to the fenestration site of ASIC3 may increase the conductance of ions through the channel by relieving the block of the central pore by native phospholipid tails.

### The spontaneous binding of DHA protects hASIC3’s central pore

Next, we verified that this mechanism can occur in situations where the lipid is not docked to the channel but needs to diffuse and reach its binding site. To determine if DHA can reach its binding site and compete with the POPC molecules that bind at or near the R63 site, DHA was randomly embedded into the lipid layer and allowed to spontaneously bind to hASIC3 in the open state (Fig. 5, Traj. 5). DHA bound to the R63 site very rapidly (Fig. S3) and a DHA occupancy heat map shows that DHA concentrated around hASIC3 during the course of the simulation (Fig. 5A). In this simulation, DHA binding to R63 was more stable than the binding of POPC likely due to the differences in head group size and charge (Fig.1A). As in the DHA docked simulations, the spontaneous binding of DHA also prevented the insertion of the POPC tail into the pore of ASIC3 (Fig. 5B). Interestingly, DHA molecules themselves could also enter the pore (Fig. 5A) but this entry was transient and unstable, with DHA molecules exiting the pore in microseconds. This transient entry obstructed the continuous water wire, leading to intermittent dehydration of the aqueous pathway (Fig.5C) and obstruction in ion permeations as well (Fig. 5D). As shown in Fig. 5C, the black region indicates the temporary absence of water molecules due to transient pore blockade by DHA. However, when DHA exited the pore, the water wire regained its continuity, allowing ions to permeate through the channel (Fig. 5D). For instance, no ion permeation events occur between 5 to 7 µs due to DHA transiently entering the pore. At 7 µs, multiple ion permeation events were observed as DHA exits the pore. These transient blockade and re-opening events are illustrated in the snapshots presented in Fig. 5E-H. Overall, the presence of DHA, whether docked to the TM helices of hASIC3 or partitioned into the lipid layer, prevented POPC lipids from accessing and penetrating the pore, ensuring high permeability for water and Na^+^ ions to flow through the channel.

**Figure 5.**
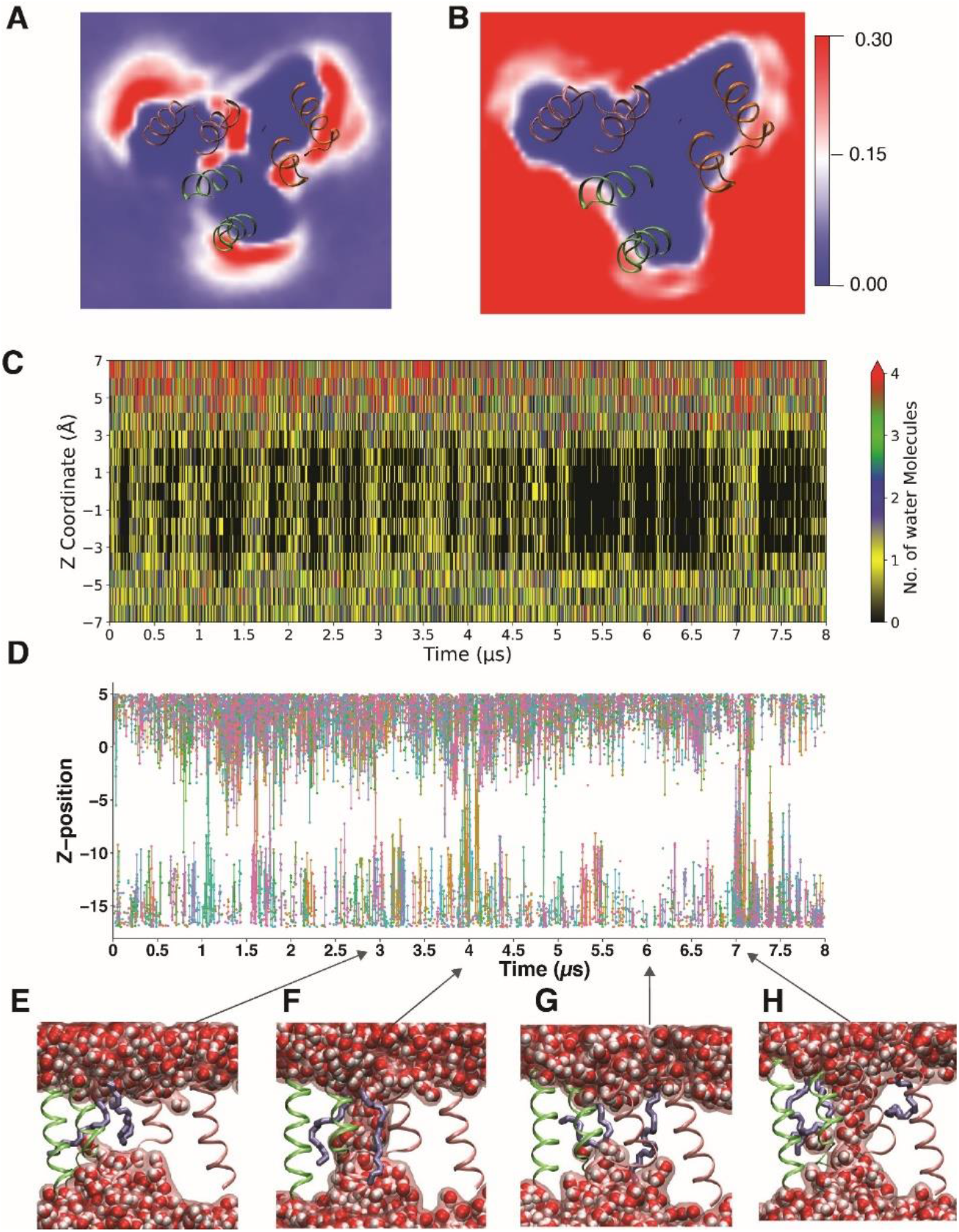
Spontaneous DHA binding to hASIC3 prevents POPC from blocking the pore. A) Occupancy volume map for DHA in Traj. 5. The color bar represents the fractional occupancy of DHA in space, where 0.3 corresponds to at least 30% occupancy during the simulation. B) Occupancy volume maps of POPC in Traj. 5. The color bar represents the fractional occupancy of POPC in space, where 0.3 corresponds to at least 30% occupancy during the simulation. DHA, randomly partitioned into the membrane, binds to R63 and prevents POPC from blocking hASIC3. **C)** Water permeation heatmap, which shows the number of water molecules along the Z-coordinate (Y-axis) over time (X-axis). The scale bar shows that anywhere from 0-4 or more water molecules can be present along the Z-coordinate of the pore region. **D)** Ion permeation analysis through the duration of Traj. 5, where each color represents a different ion permeation event. **E)** Molecular image, from Traj. 5, visualizing two DHA (ice blue licorice) molecules blocking continuous water flow through the central pore at 3µs. **F)** Molecular image at 4 µs visualizing the continuous water wire. **G & H)** Snapshots at 6 and 7 µs respectively showing the transient block of water flow by DHA and block relief in H showing continuous water wire.

### Single-channel current amplitudes are consistent with a fast block mechanism

Finally, to experimentally measure changes in the ion conductance of ASIC3 in the absence and presence of potentiating lipids, we utilized single-channel patch-clamp recordings to assess the unitary conductance of individual opening events. Single-channel patch-clamp measurements provide a way to analyze the mechanisms of ion channel block. If we treat the block of ASIC3 conduction by POPC lipid tails as a fast block of current, the classical description of open-channel block in single-channel current measurements predicts that a very fast block results in a measured decrease in the single-channel current amplitude. Thus, we would predict that if the POPC tail block is fast, then the addition of a potentiating lipid (NAAAs or PUFAs), would result in relief of this fast block and an overall increase in the single channel current amplitude.

To test this, we recorded unitary currents from ASIC3 channels present in outside-out patches excised from transiently transfected CHO cells. Channel activity was measured in response to rapid switches from pH 8 to pH 5.5 at a voltage of -80mV as we did in our whole cell measurements. Representative currents from a patch containing 1 channel can be seen in Figure 6A. From this recording, we created an all-point histogram of current amplitude and fit with two Gaussians which shows two populations of current amplitudes, one for open channels and one for closed (Fig. 6A right). We repeated these experiments after preincubating our cells in 10μM DHA for 10 minutes. A representative patch with two channels can be seen in Figure 6B. Again, we used an all-points histogram fit with two Gaussians to determine the single-channel current amplitude (Fig. 6B, right). For comparison, we only show the current levels corresponding to the closed state and the single opening event and have not displaced the region of the histogram corresponding to the two-channel level of current. We performed this analysis on 5 patches for each condition. The result of this analysis showed that the single channel current amplitude for ASIC3 channels increased from -1.5 +/-0.02 pA to -1.66 +/-0.01pA in the presence of 10μM DHA (Fig. 6C), resulting in a 10.7% increase in the unitary channel conductance. These data are consistent with the observed phenomenon in our simulations where DHA relieves the fast block of POPC tails by displacing them from the pore of ASIC3.

**Figure 6.**
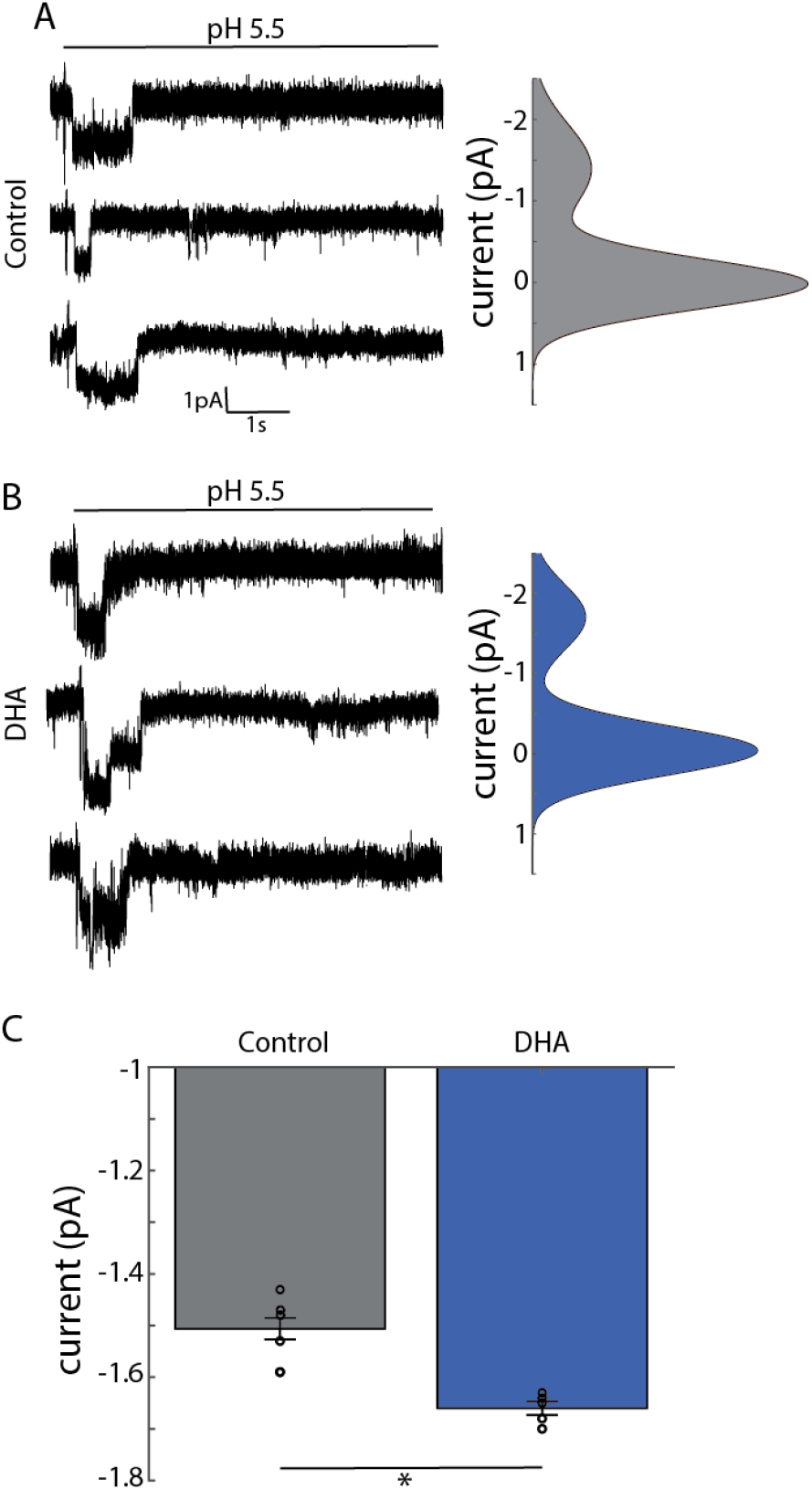
DHA increases single-channel current amplitude in ASIC3. **A)** Representative traces of single-channel recordings under control conditions showing three sweeps from a patch with a single rASIC3 channel elicited by a switch to pH 5.5. (left). Gaussian fits all-points histogram of current amplitudes (right), which comprised of all sweeps from this patch, were plotted to fit peaks in the data. **B)** Representative traces of single-channel recordings under DHA conditions showing three sweeps from a patch with a single rASIC3 channel elicited by a switch to pH 5.5 in the presence of 10µM DHA (left). Gaussian fits all-points histogram of current amplitudes (right), which comprised of all sweeps from this patch, were plotted to fit peaks in the data. **C)** Bar graph showing the average single-channel current amplitude +/-SEM from patches in the presence and absence of 10µM DHA. Statistical comparison was made using an unpaired T-test. (n=5 patches for control and n=5 patches for DHA, ^*^=p<2*10^−5^).

## Discussion

### Structural Basis for PUFA-Mediated Enhancement of ASIC3 Currents

ASIC3 is sensitive to regulation by a number of single acyl chain lipids from various classes including PUFAs, NAAAs, and lysophosphatidic cholines (LPCs) (14,18,47).

Direct binding of these lipids causes multiple changes in channel gating, all of which lead to an increase in ASIC3 current. This potentiation can even cause channel activation in the absence of acidification. One of these gating changes produces an increase in the maximal current even at saturating proton concentrations. Here, we suggest a potential explanation for this change in channel function using MD simulations and patch-clamp electrophysiology. We hypothesize that under control conditions, the pore of ASIC3 undergoes rapid transient block from the tails of the native phospholipids. This block reduces the single-channel conductance of the channel. Potentiating lipids, like AG or DHA, can then outcompete the native phospholipids for binding sites on the channel that allow for tail access to the fenestrations, reducing fast block by the POPC tails and increasing the single-channel current amplitude.

This competition between PUFA and POPC occurs at a key arginine residue (R63) that is required for DHA regulation in ASIC3. The polar head groups of both PUFAs and POPC molecules are composed of negatively charged groups, facilitating their binding to positively charged protein residues. Nonetheless, the binding of PUFAs is significantly more stable than that of POPC. Notably, while both lipid molecules feature negatively charged oxygen atoms in their head groups, POPC additionally contains a positively-charged and bulky choline group in its head group. This structural disparity alters the charge distribution within the lipid molecules and influences their interactions with the binding residues. Hence, the strength of the ionic bond between POPC and the arginine residue is lower than that between PUFAs and the same residue, allowing PUFAs to act as gatekeepers preventing POPCs from the lipid layer from reaching the pore. When PUFAs are absent from the fenestration site, these POPCs can then access the central pore, transiently and frequently impeding both water and ion permeation.

### Interactions between lipids and channel fenestrations may be common

This idea of lipid tail occlusion of ion channels has been suggested through multiple studies. *In silico* work on a number of voltage-gated sodium channel isoforms shows that lipid tails often penetrate the fenestrations in these channels, suggesting a hydrophobic nature of these pathways (48). Inward-rectifier (Kir) potassium channels are thought to be partially gated by a mechanism similar to the one we describe here whereby anionic lipid head groups bind tightly to the channel at the intracellular surface and allow for the tails to enter fenestrations in the channel and occlude ion flow (49). In TREK1 channels, POPA lipid tails interact with critical residues near the selectivity filter of the channel near the permeation path of the channel (50).

While our data here provide a possible explanation for how I_Max_ changes upon single acyl chain lipid binding to ASIC3, more work is needed to understand how this binding alters both the rate of desensitization and the sensitivity of the channel to proton concentration. It is possible that this tail-block mechanism could impact all of the gating changes measured in ASIC3 upon the addition of single acyl chain lipids. Displacement of the occluding phospholipid tail would likely increase channel open times which could lead to a slowing of the rate of channel desensitization and could result in an increased overall activity at a more neutral pH. This would suggest that the lipid tail is acting like a gate for the channel similar to a proposed mechanism for Kir gating (49). However, more work is needed to determine if this is the case.

The binding site for several small molecule inhibitors of ASICs resides in the pore region near the bottom of the fenestrations of the channel in the region where we observe the lipid tail (11,51). The lipid composition of the membrane has been shown to alter the pharmacological activity of many existing drug molecules on other ion channels, In CaV1.1, a lipid tail that putatively blocks the channel fenestration has been resolved in cryoEM structures(52). This tail was suggested to help keep ligands that bind in the pore region trapped in their binding sites. In sodium channels, access to the drug binding site, especially for hydrophobic blockers, depends on the size and dynamics of the fenestrations as well (53). It will be interesting to determine if the binding of pore blockers in ASICs depends on this fenestration as well as the nature of the lipid membrane.

### Limitations and model for the change in I_Max_

This hypothesis was derived using *in silico* models but is challenging to prove experimentally. Our single-channel measurements show that DHA increases the single-channel current amplitude of ASIC3, which is consistent with the idea that these lipids may relieve a fast block caused by the presence of POPC lipid tails in the pore. However, it is certainly possible that the change in current amplitude arises from an alternative mechanism. The binding of lipids may allosterically alter the conduction pathway changing the pathway through which ions flow. Beyond that, our simulations are based on a homology model that was derived from the only available open-state structure of an ASIC. This structure was a toxin-bound open state of ASIC1 from chicken with a wider than expected pore that was also missing a key piece of the permeation path: the n-terminal re-entrant loop that is now thought to impact permeation and selectivity of the channel (54).

Despite these limitations, we propose a model whereby the phospholipids in the extracellular leaflet of the plasma membrane can transiently bind to an arginine on TM1 of ASIC3 positioning tail groups to penetrate the fenestrations in the transmembrane segments of the channel. In doing so, these lipid tails occlude the permeation pathway of the channel and reduce the conductance of the channel. Single acyl chain lipids such as PUFAs and NAAAs can displace the POPC from this region relieving the tail block of the channel and increasing the channel conductance. Future work will need to determine if ASIC3 conductance depends on the composition of the outer leaflet of the membrane. Collectively, our findings put forth a previously unobserved but potentially widespread mechanism through which PUFAs may regulate ion channel activity.

## Author contributions

All authors contributed to conception and design.

R.B. and J.L. carried out MD simulations and trajectory analysis.

R.R., R.K., and J.B. performed electrophysiology measurements and analysis. writing-original draft: R.B., J.B., and J.L.

Writing-review & editing: R.B., R.R., J.B., and J.L.

## Competing interests

The authors declare that they have no competing interests.

## Acknowledgements

Computer resources came from a Maximize ACCESS allocation through project BIO210015, an allocation (MCB200085P) on Antons at the Pittsburgh Supercomputing Center provided by the National Center for Multiscale Modeling of Biological Systems through National Institutes of Health grant P41GM103712-1 and from a loan from D. E. Shaw Research, and a Frontera Pathways allocation (MCB21012) at the Texas Advanced Computing Center (TACC). Research reported in this publication was supported by an Institutional Development Award (IDeA) from the National Institute of General Medical Sciences of the National Institutes of Health under award number P20GM130460 to JL as well as R35GM137912 from the National Institute of General Medical Sciences to JB.

## Notes

### Competing Interest Statement

The authors have declared no competing interest.

