## Supplementary file for "Molecular Insights into Single Chain Lipid Modulation of Acid-Sensing Ion Channel 3"

This file includes:

Figures S1 to S3
Videos S4 and S5

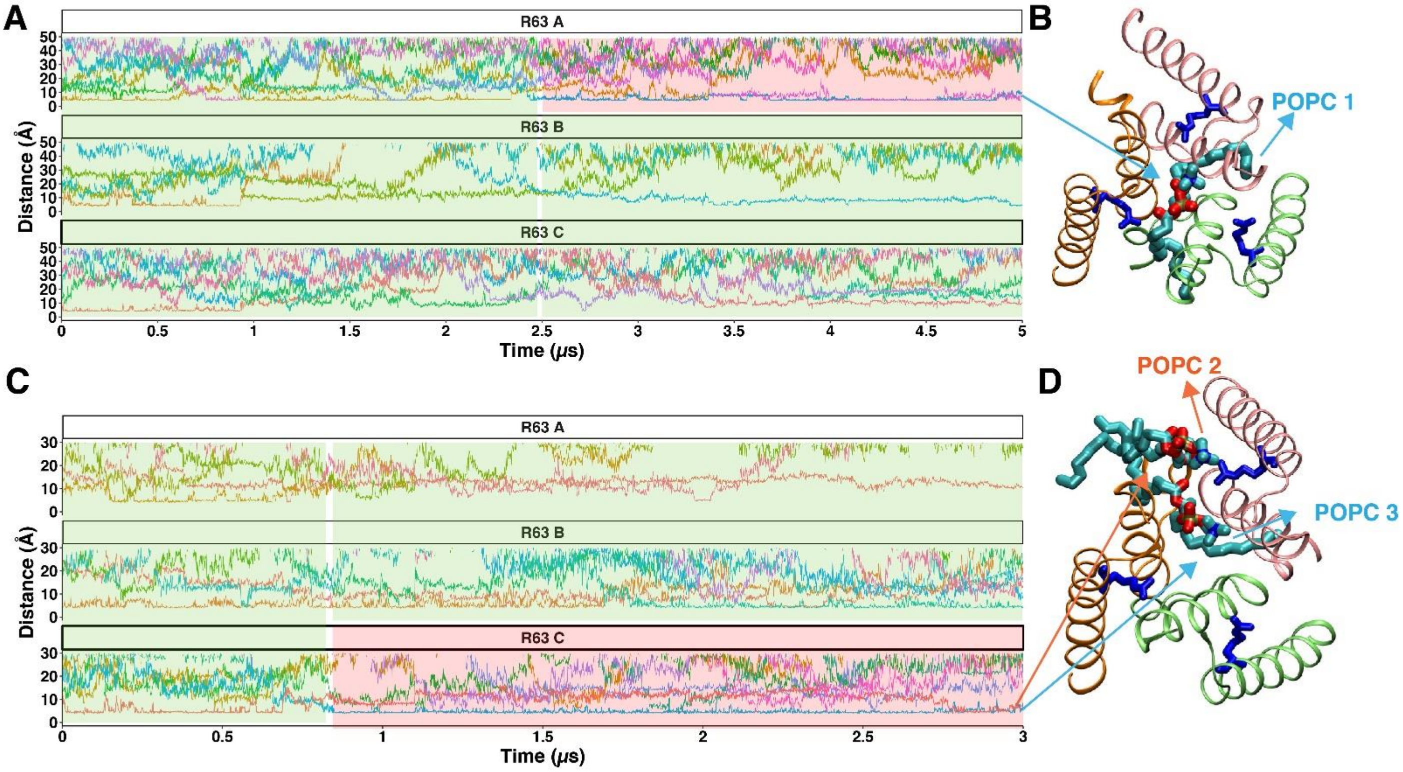


**Fig. S1: POPC bound to R63 blocks the pore of open-state hASIC3: A)** Distance plot of POPCs to three R63 in different chains of hASIC3 in the PUFA-free simulation (Traj.1). Different colored traces represent different POPC molecules. The green-highlighted region indicates where POPC molecules are bound to R63 but do not access the pore. The red-highlighted region indicates where the POPC bound to R63 enters and blocks the pore. **B)** Molecular image of the last frame of Traj.1, showing POPC (represented in cyan licorice) bound to R63 (represented in blue licorice) blocking the pore. **C)** In the same manner to Fig. S1A, plot depicts the distance between POPCs and R63 in Traj.2. Two POPC molecules bound to R63 block the pore. **D)** Molecular image of the last frame of Traj.2, showing two POPC molecules (cyan licorice) bound to R63 (blue licorice) blocking the pore.

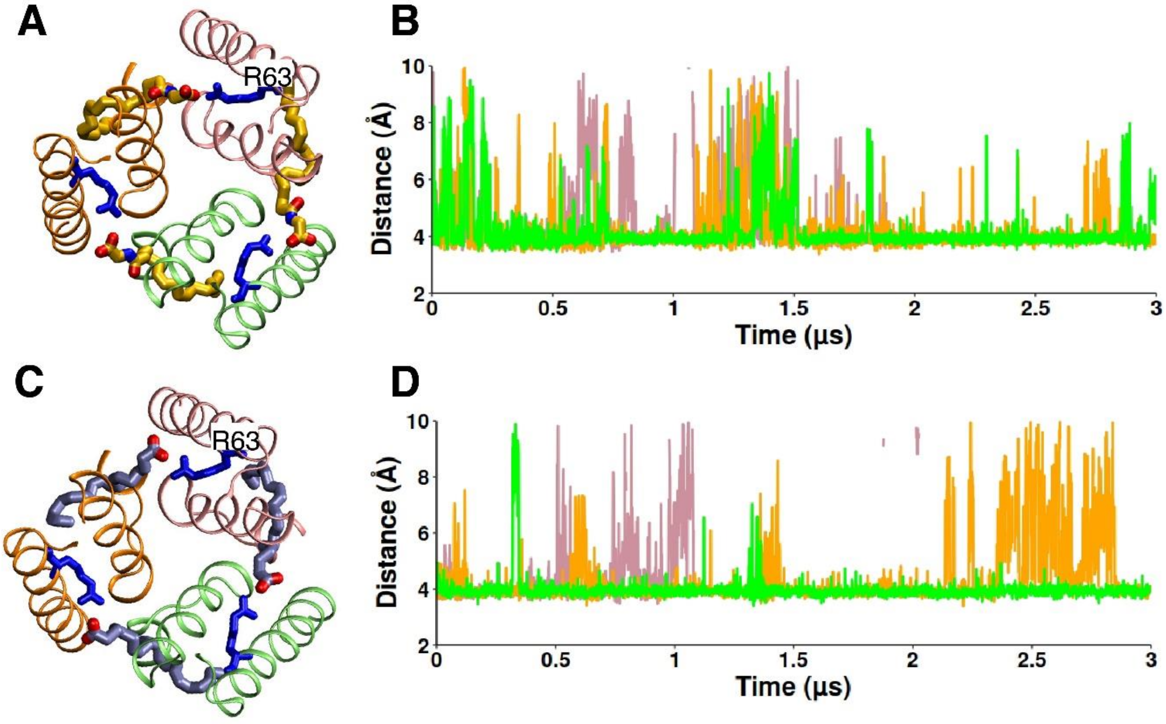


**Fig. S2: Stable binding of AG and DHA to R63: A)** Molecular image visualizing the initial frame of Traj.3, showing 3 AG molecules (represented in yellow licorice) bound to R63 (blue licorice) of 3 chains of hASIC3. **B)** Distance plot between the carboxyl carbon of AG and $C_Z$ of R63 of the corresponding chain throughout the simulation, with three colors representing the AG bound to R63 of three chains. **C)** Molecular image visualizing the initial frame of Traj.4, showing 3 DHA molecules (represented in ice blue licorice) bound to R63 (blue licorice) of 3 chains of hASIC3. **D)** Distance plot of DHA and R63 in the same manner to Fig. S2B.

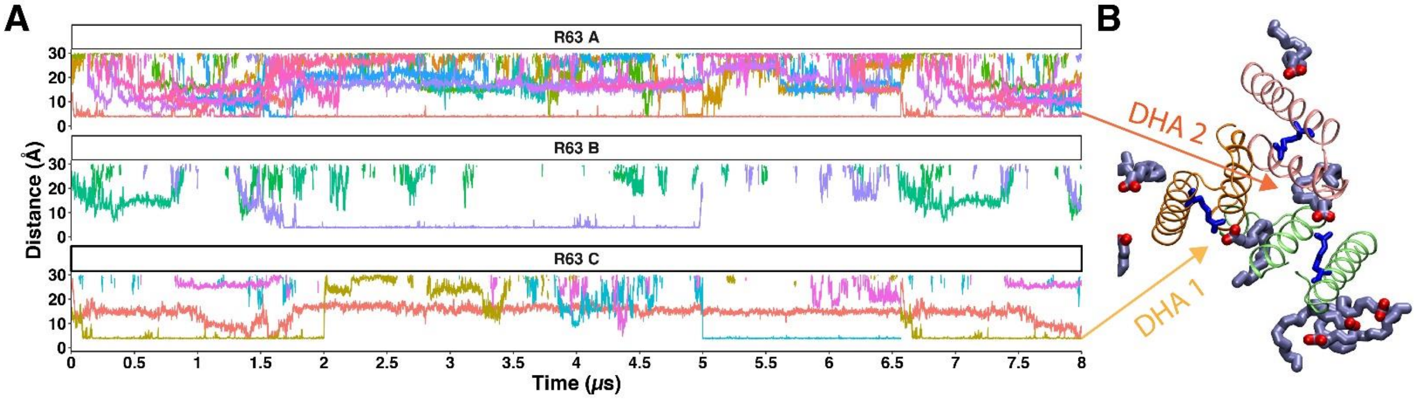


**Fig. S3: Spontaneous binding of DHA to R63 (Traj. 5)**: **A)** Distance plot illustrating the proximity between carboxyl carbon of DHA and $C_Z$ of R63 in three chains, with different colored traces indicating distinct DHA molecules. Spontaneous binding is observed within first few nanoseconds of simulation. **B)** Snapshot of last frame of traj.5 depicting 2 DHA molecules (iceblue licorice) bound to R63 (represented by blue licorice).

**Video S4**: Top view of hASIC3 pore region of PUFA-free simulation (Traj.1) showing POPC (represented in cyan licorice) accessing the pore through fenestration site and blocking the pore. Three chains are represented in three different colors.

**Video S5**: Top view of the hASIC3 pore region from the 3-AG bound simulation (Traj.3), showing AG (represented in yellow licorice) bound to R63, preventing POPC from accessing and blocking the pore.
